# Negative short-range genomic autocorrelation of causal effects on human complex traits

**DOI:** 10.1101/2020.09.23.310748

**Authors:** Armin P. Schoech, Omer Weissbrod, Luke J. O’Connor, Nick Patterson, Huwenbo Shi, Yakir Reshef, Alkes L. Price

## Abstract

Most models of complex trait genetic architecture assume that signed causal effect sizes of each SNP (defined with respect to the minor allele) are uncorrelated with those of nearby SNPs, but it is currently unknown whether this is the case. We develop a new method, autocorrelation LD regression (ACLR), for estimating the genome-wide autocorrelation of causal minor allele effect sizes as a function of genomic distance. Our method estimates these autocorrelations by regressing the products of summary statistics on distance-dependent LD scores. We determined that ACLR robustly assesses the presence or absence of nonzero autocorrelation, producing unbiased estimates with well-calibrated standard errors in null simulations regardless of genetic architecture; if true autocorrelation is nonzero, ACLR correctly detects its sign, although estimates of the autocorrelation magnitude are susceptible to bias in cases of certain genetic architectures. We applied ACLR to 31 diseases and complex traits from the UK Biobank (average *N*=331K), meta-analyzing results across traits. We determined that autocorrelations were significantly negative at distances of 1-50bp (*P* = 8 × 10^−6^, point estimate −0.35 ±0.08) and 50-100bp (*P* = 2 × 10^−3^, point estimate −0.33 ± 0.11). We show that the autocorrelation is primarily driven by pairs of SNPs in positive LD, which is consistent with the expectation that linked SNPs with opposite effects are less impacted by natural selection. Our findings suggest that this mechanism broadly affects complex trait genetic architectures, and we discuss implications for association mapping, heritability estimation, and genetic risk prediction.

## Introduction

Analyses of complex trait genetic architecture, i.e. the genome-wide distribution of causal genetic effects, have substantially improved our understanding of complex trait biology, informing genetic association studies and genetic risk prediction^1–16^. One aspect of genetic architecture that has not been well-studied is genomic autocorrelation, i.e. autocorrelation of signed causal effect sizes (here defined with respect to the minor allele) as a function of genomic distance. On the one hand, genomic autocorrelation could potentially be positive, if the minor alleles of two nearby SNPs disrupt the same functional element^17;18^. On the other hand, genomic autocorrelation could potentially be negative, if linked SNPs with opposite trait effects escape negative selection, a phenomenon known as linkage masking^19^. Several previous studies have provided specific examples of concordant^20^ and opposite^19;21^ trait effects of nearby SNPs in humans, as well as concordant^22^ and opposite^22;23^ trait effects in model organisms. However, genomewide genomic autocorrelation has not yet been investigated. It is therefore unclear if genome-wide autocorrelation exists, and if so, what its sign, magnitude, and genomic distance range is.

Here, we introduce a new method, autocorrelation LD regression (ACLR), for estimating the genomewide autocorrelation of causal genetic effects as a function of genomic distance. Specifically, we define genomic autocorrelation as the genome-wide Pearson correlation coefficient between causal minor allele effects of all pairs of genetic variants that are a specific base pair distance apart. ACLR regresses products of genome-wide association study (GWAS) summary statistics on distance-dependent linkage disequilibrium (LD) scores. We evaluate ACLR by performing simulations under a range of genetic architectures, and apply ACLR to 31 disease and complex traits from the UK Biobank^24^ (average *N*=331K). We further investigate how genomic autocorrelations of nearby SNPs depend on their pairwise LD. We discuss the implications of genomic autocorrelation for analyses of heritability and genome-wide association studies.

## Results

### Overview of methods

ACLR estimates the genome-wide autocovariance and autocorrelation of causal minor allele effects as a function of base pair distance by regressing products of GWAS summary statistics on distance-dependent LD scores. Products of GWAS summary statistic of two SNPs will tend to be positive if their causal effects are correlated, but also if both SNPs are in LD with (i.e. tag) the same causal SNP. Intuitively, ACLR accounts for the effect of tagging to specifically infer the correlation of causal effects of SNP pairs depending on the base pair distance between them.

In detail, ACLR partitions SNP pairs into several base pair distance ranges (e.g. 1-50bp, 50-100bp, etc.) and assumes a fixed causal effect covariance of per-allele effect sizes for each distance range. We show that the product of GWAS summary statistics of two SNPs *i* and *j* has the following expectation (see Methods):

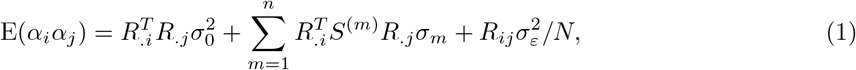

where *R* is the LD matrix (specifically the minor allele count covariance matrix), 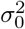 is the genomewide causal per-allele effect size variance, *S*^(*m*)^ is the design matrix of distance range *m* (with 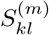 taking values of 0 or 1 indicating whether the distance between SNP *k* and SNP *l* falls into the *m*^th^ distance range), *σ_m_* is the genome-wide causal minor allele effect size autocovariance for distance range *m*, 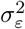 is the variance of environmental (non-genetic) effects on the trait, and *N* is the sample size. *σ_m_* can hence be inferred by regressing *α_i_α_j_* on distance-dependent LD scores 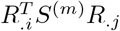. Unlike previous methods that use external reference LD panels to estimate LD between SNPs^5;25;26^, we use in-sample LD estimates from imputed genotypes of the target samples (see Methods), which has been preferred in some recent analyses^27–29^. Distance-dependent causal minor allele effect autocorrelation estimates 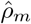 are computed by dividing the corresponding autocovariance estimates 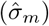 by estimates of the genome-wide causal perallele effect size variance 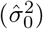. Standard errors are estimated via genomic block-jackknife (see Methods). We emphasize that autocovariances and autocorrelations are reported with respect to per-allele effects, and not per-standardized allele effects.

Systematic differences in causal per-allele effect size variances between SNPs with different minor allele frequencies (MAF) and levels of LD can lead to substantial bias in related approaches that estimate different quantities^4;8;30–32^. Thus, ACLR allows for “MAF- and LD-corrected” estimates that explicitly allow for these differences (see Methods). Specifically, we model different causal per-allele effect size variances using 10 common SNP MAF bins, 10 low-frequency SNP MAF bins, and 6 LD-related annotations^8;33^. We have publicly released open-source software implementing the ACLR method (see URLs).

### Simulations

We evaluated ACLR in simulations under a range of genetic architectures using UK Biobank genotype data (see Methods). We used the same set of 337,502 unrelated British-ancestry individuals as in our analyses of UK Biobank traits. Due to the large number of simulations, we restricted most simulations to chromosome 22 (232,046 SNPs with MAF ≥ 0.1%); however, we also performed a selected set of simulations using genome-wide data (16,525,661 SNPs with MAF ≥ 0.1%). We performed simulations with uncorrelated causal minor allele effects (null simulations), as well as simulation with nonzero genomewide autocorrelations (causal simulations). ACLR assumes that causal effect autocovariances are constant for SNP pairs within each distance range, but we included causal simulations in which this assumption is violated. We set genome-wide heritability to either 0.5 or 0.2. In most simulations we specified standardized causal effect sizes using the baseline-LF model of Gazal et al.^33^, which models MAF- and LD-dependent effects; however, we also performed simulations in which causal per-allele effect sizes are independent of MAF and LD. We evaluated the accuracy of both autocovariance and autocorrelation estimates, as autocovariance estimates were more accurate (see below) but autocorrelation estimates are more intuitively interpretable.

We first performed null simulations to assess bias and calibration when true causal effects are uncorrelated. Results are reported in Table 1. We determined that MAF- and LD-corrected autocovariance and autocorrelation estimates were unbiased, with well-calibrated standard errors, for all genetic architectures simulated. However, as expected, estimates without MAF- and LD-correction were biased in the case of MAF- and LD-dependent genetic architectures. Although these simulations used data from chromosome 22 only, a subset of simulations using genome-wide data yielded similar conclusions (see Supplementary Table 1).

**Table 1:** ACLR results in null simulations with zero autocorrelation

| MAF-LD<br>corrected | MAF-LD<br>architect | $h^2$ | $\hat{\sigma}_{1-100bp}$ | $\hat{\rho}_{1-100bp}$ | $\hat{\sigma}_{100-1000bp}$ | $\hat{\rho}_{100-1000bp}$ |
| --- | --- | --- | --- | --- | --- | --- |
| yes | yes | 0.5 | $-0.008 \pm 0.008$<br>(0.008) | $-0.004 \pm 0.010$<br>(0.011) | $0.002 \pm 0.001$<br>(0.001) | $0.002 \pm 0.002$<br>(0.002) |
| yes | no | 0.5 | $-0.002 \pm 0.011$<br>(0.011) | $0.010 \pm 0.011$<br>(0.011) | $-0.001 \pm 0.002$<br>(0.002) | $-0.001 \pm 0.002$<br>(0.002) |
| yes | yes | 0.2 | $-0.007 \pm 0.008$<br>(0.008) | $-0.003 \pm 0.010$<br>(0.010) | $0.002 \pm 0.001$<br>(0.001) | $0.002 \pm 0.002$<br>(0.002) |
| no | no | 0.5 | $-0.003 \pm 0.011$<br>(0.011) | $0.013 \pm 0.011$<br>(0.011) | $-0.005 \pm 0.002$<br>(0.002) | $-0.004 \pm 0.002$<br>(0.002) |
| no | yes | 0.5 | $-0.182 \pm 0.008$<br>(0.009) | $-0.212 \pm 0.010$<br>(0.011) | $-0.052 \pm 0.001$<br>(0.001) | $-0.063 \pm 0.002$<br>(0.002) |

For each genetic architecture simulated, we report autocovariance estimates ( $\hat{\sigma}$ ) and autocorrelation estimates ( $\hat{\rho}$ ) for 1-100bp and 100-1000bp genomic distance ranges, and their empirical standard errors. We also report mean jackknife standard error estimates in parentheses. Autocovariance estimates ( $\hat{\sigma}$ ) are scaled based on a true variance of 1 for ease of comparison; true values of $\sigma$ and $\rho$ are equal to 0. "MAF-LD corrected" denotes MAF- and LD-corrected estimates. "MAF-LD architect" denotes MAF- and LD-dependent generative architectures. $h^2$ denotes the heritability simulated. Results are averaged across 1000 simulations in each case.

We then performed causal simulations, to assess how well ACLR can detect and quantify nonzero autocovariance and autocorrelation. We first discuss autocovariance estimates in causal simulations (Table 2). We determined that MAF- and LD-corrected autocovariance estimates were unbiased, with well-calibrated standard errors, for all genetic architectures in which true causal effect autocovariances were constant for SNP pairs within each distance range (as assumed by the inference method). On the other hand, autocovariance estimates were biased when true causal effect autocovariances were not constant for SNP pairs within each distance range - although the sign of the nonzero autocovariance was still correctly inferred. This bias due to unmodeled variation in autocovariance signal across pairs of SNPs is analogous to biases that can arise in heritability estimation due to unmodeled variation in per-SNP heritability^4;8;30–32^.

**Table 2:** ACLR results in causal simulations with nonzero autocorrelation

| MAF-LD<br>architect | $h^2$ | Const<br>covar | $\sigma_{1-100bp}$ | $\hat{\sigma}_{1-100bp}$ | $\hat{\rho}_{1-100bp}$ | $\sigma_{100-1000bp}$ | $\hat{\sigma}_{100-1000bp}$ | $\hat{\rho}_{100-1000bp}$ |
| --- | --- | --- | --- | --- | --- | --- | --- | --- |
| no | 0.5 | yes | 0.200 | $0.203 \pm 0.016$<br>(0.017) | $0.215 \pm 0.016$<br>(0.017) | 0.000 | $-0.000 \pm 0.003$<br>(0.003) | $0.003 \pm 0.003$<br>(0.003) |
| no | 0.5 | yes | -0.200 | $-0.206 \pm 0.014$<br>(0.015) | $-0.197 \pm 0.014$<br>(0.014) | 0.000 | $-0.002 \pm 0.003$<br>(0.003) | $0.000 \pm 0.003$<br>(0.003) |
| yes | 0.5 | yes | 0.230 | $0.220 \pm 0.014$<br>(0.014) | $0.297 \pm 0.018$<br>(0.018) | 0.000 | $-0.005 \pm 0.003$<br>(0.003) | $-0.002 \pm 0.003$<br>(0.003) |
| yes | 0.5 | yes | -0.250 | $-0.256 \pm 0.009$<br>(0.009) | $-0.356 \pm 0.013$<br>(0.014) | 0.000 | $-0.002 \pm 0.002$<br>(0.002) | $-0.000 \pm 0.003$<br>(0.003) |
| yes | 0.2 | yes | 0.230 | $0.219 \pm 0.014$<br>(0.015) | $0.296 \pm 0.018$<br>(0.018) | 0.000 | $-0.005 \pm 0.003$<br>(0.003) | $-0.002 \pm 0.003$<br>(0.003) |
| yes | 0.2 | yes | -0.250 | $-0.255 \pm 0.009$<br>(0.009) | $-0.354 \pm 0.013$<br>(0.014) | 0.000 | $-0.002 \pm 0.002$<br>(0.002) | $-0.000 \pm 0.003$<br>(0.003) |
| yes | 0.5 | no | 0.149 | $0.034 \pm 0.009$<br>(0.009) | $0.051 \pm 0.010$<br>(0.010) | 0.033 | $0.009 \pm 0.001$<br>(0.001) | $0.010 \pm 0.002$<br>(0.002) |
| yes | 0.5 | no | -0.149 | $-0.048 \pm 0.007$<br>(0.007) | $-0.056 \pm 0.009$<br>(0.009) | -0.033 | $-0.007 \pm 0.001$<br>(0.001) | $-0.009 \pm 0.001$<br>(0.001) |

For each genetic architecture simulated, we report the true autocovariances ( $\sigma$ ), autocovariance estimates ( $\hat{\sigma}$ ) and autocorrelation estimates ( $\hat{\rho}$ ) for 1-100bp and 100-1000bp genomic distance ranges, and their empirical standard errors. We also report mean jackknife standard error estimates in parentheses. True autocovariances ( $\sigma$ ) and their estimates ( $\hat{\sigma}$ ) are scaled based on a true variance of 1 for ease of comparison; thus, the true $\rho$ is equal to the true $\sigma$ . All estimates were MAF- and LD-corrected. "MAF-LD architect" denotes MAF- and LD-dependent generative architectures. $h^2$ denotes the heritability simulated. "Const covar" denotes genetic architectures in which true autocovariances ( $\sigma$ ) are constant for SNP pairs within each distance range. Results are averaged across 1000 simulations in each case.

We next discuss autocorrelation estimates in causal simulations (Table 2). In simulations without MAF- and LD-dependent genetic architectures, autocorrelation estimates were unbiased, with well-calibrated standard errors, analogous to null simulations. However, in simulations with MAF- and LD-dependent genetic architectures, autocorrelation estimates were biased - even when true causal effect autocovariances were constant for SNP pairs within each distance range, such that autocovariance estimates were unbiased; in each case, the sign of the nonzero autocorrelation was still correctly inferred. Since autocorrelation estimates are computed by dividing autocovariance estimates by SNP causal effect variance estimates, bias in autocorrelation estimates in the absence of bias in autocovariance estimates can be explained by bias as well as noise in SNP causal effect variance estimates, although we did not observe the latter playing a significant role in our simulation results.

In summary, ACLR yields unbiased autocovariance and autocorrelation estimates with well-calibrated standard errors in all null simulations. ACLR detects nonzero autocovariance and autocorrelation and correctly infers their sign in all causal simulations, although the magnitude of the autocovariance and autocorrelation estimates can be biased depending on the genetic architecture simulated. Thus, any significantly nonzero autocovariance/autocorrelation reported by ACLR is a robust result, in the sense that the true autocovariance/autocorrelation will be nonzero with the reported sign.

### Negative short-range genomic autocorrelation across 31 UK Biobank traits

We estimated the genome-wide autocovariance and autocorrelation of causal minor allele effect sizes, as a function of genomic distance, for 31 strongly heritable diseases and complex traits from the UK Biobank (Supplementary Table 2; average N=330,618). We analyzed 16, 525, 661 genome-wide SNPs with MAF ≥ 0.1% and applied ACLR to summary association statistics computed using BOLT-LMM^7;34^ together with in-sample LD from imputed genotypes (see Methods). We meta-analyzed results across 25 independent traits with pairwise genetic correlation less than 0.5 (see Methods).

Autocorrelation estimates are reported in Figure 1 and Supplementary Table 3, and the corresponding autocovariance estimates are reported in Supplementary Table 4 (autocovariance estimates are less intuitively interpretable, with absolute values ranging from the order of 10^−8^ to 10 ^−10^ across traits for variants less than 1000bp apart). We detected significantly negative autocorrelations in the meta-analysis at distances of 1-50bp (*P* = 8 × 10^−6^, point estimate −0.35±0.08) and 50-100bp (*P* = 2 × 10^−3^, point estimate −0.33±0.11). Results were fairly consistent across traits (e.g. 1-50bp: negative estimates for 15/16 traits with standard errors < 0.5), and we did not detect statistically significant heterogeneity across traits for any distance range (see Supplementary Table 5). We observed no significant deviation from zero for 100-200bp and 200-500bp, and a slightly positive autocorrelation at 500-1000bp (*P* = 3 × 10^−3^, point estimate 0.07 ± 0.02). No single-trait autocorrelation estimate was significantly different from zero for any distance range after Bonferroni correction, though several were nominally significant at a significance threshold of 0.05. We also estimated the effect autocorrelation for SNPs more than 1000bp apart, but estimates were very small (absolute value < 10^−3^ for all traits) and not significantly different from zero (Supplementary Table 6).

**Figure 1:**
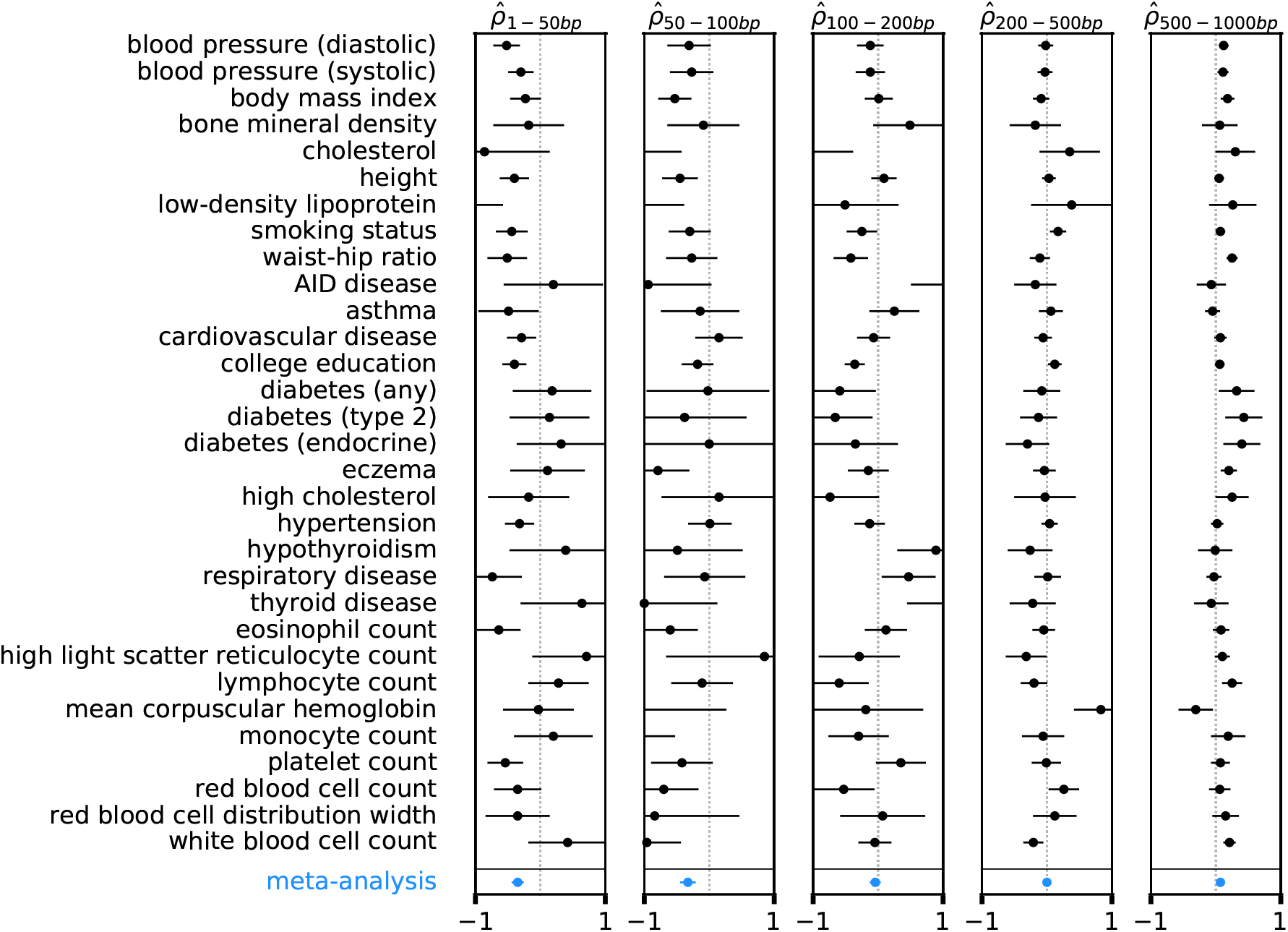
ACLR results for 31 UK Biobank traits. We report autocorrelation estimates 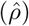 across 5 different genomic distance ranges for each of 31 UK Biobank traits, including 9 quantitative traits, 13 case-control traits, as well as 9 quantitative blood count traits. We also show meta-analysis results across a subset of 25 independent traits in the last row. Error bars denote block-jackknife standard errors. Numerical results and autocovariance estimates are reported in Supplementary Tables 3 and 4 respectively.

In a more stringent meta-analysis across 13 strictly independent traits with pairwise genetic correlation less than 0.1 (Supplementary Table 7), the negative autocorrelation at 1-50p remained highly significant (*P* = 10^−4^) but the remaining autocorrelations were not significant after correcting for hypotheses tested for six distance ranges. We thus consider the negative short-range autocorrelation to be a robust result. As noted above, our simulations showed that conclusions about deviations from zero are robust, as ACLR was unbiased and well-calibrated in null simulations regardless of genetic architecture. Our simulations also showed that, in the case of true nonzero autocorrelations, autocorrelation estimates can have biased magnitudes. Specifically, autocorrelation estimates can be biased if true causal effect autocovariances are not constant for SNP pairs within each distance range (in which case autocovariance estimates are biased); it is possible that this phenomenon impacts our results on real traits. Autocorrelation estimates can also be biased when true causal effect autocovariances are constant for SNP pairs within each distance range (in which case autocovariance estimates are unbiased, but autocorrelation estimates can still be biased due to noise (or bias) in per-allele effect variance estimates); it is unlikely that this phenomenon significantly impacts our results on real traits, because heritability estimates produced by ACLR based on effect variance estimates are similar to previously published estimates (Supplementary Table 2) and because we only included traits that are strongly heritable (z-score ≥ 6 for nonzero heritability; see Methods).

Previous studies have pointed out that linked SNPs with opposite effects escape the action of negative selection, a phenomenon known as linkage masking^19;23^; specifically, a haplotype harboring two SNPs with opposite effects will be less impacted by negative selection because the effects will (partially) cancel out in any individual carrying that haplotype. Linkage masking could potentially explain the systematically negative short-range genomic autocorrelation that we inferred in this study. The very short genomic distance range (on the order of 100bp) of the autocorrelation that we detected does not rule out this explanation: although substantial LD is common at much longer ranges, it is weaker at longer ranges, which could potentially reduce the effects of linkage masking^35^. In addition, linkage masking effects at very short genomic distance range could be amplified by functional similarity at this range, e.g. due to compensatory mutations in binding sites or coding regions; SNP with opposite effects may be much less likely to arise at longer ranges.

We hypothesized that, under the linkage masking hypothesis, the negative short-range autocorrelation would be stronger for pairs of SNPs with positive LD (between minor alleles). We repeated our analyses while stratifying pairs of SNPs by their pairwise LD, in addition to genomic distance (see Methods); we confirmed that this approach produces robust results in simulations (Supplementary Table 8). Consistent with our hypothesis, short-range (1-100bp) autocorrelations were significantly negative for pairs of SNPs in positive LD, but significantly positive for pairs of SNPs in negative LD, in a meta-analysis across the 25 independent traits (Supplementary Table 9). We note that positive autocorrelation of trait effects of alleles in negative LD is also consistent with linkage masking.

Heritability estimation methods commonly assume that SNP-heritability is equal to the genomewide sum of standardized per-allele effect variances^2–6;8;10;12–15^. While this assumption is correct in commonly used complex trait models that assume uncorrelated per-allele effects, it is incorrect if per-allele effects are significantly correlated, as we have demonstrated. Specifically, if linked SNPs have negatively correlated effects, the true SNP-heritability will be lower than the sum of standardized per-allele effect variances. We introduce a measure that we call “heritability shrinkage” that quantifies the relative decrease in heritability due to negatively correlated SNP effects (see Methods). Based on our autocorrelation estimates stratified by pairwise LD between pairs of SNPs (Supplementary Table 9), we derived a hypothetical heritability shrinkage estimate of (13 ± 2)% (see Methods), indicating that genomic autocorrelation may substantially impact the genetic architecture of a complex trait. We caution that our heritability shrinkage estimate should be viewed only as a hypothetical estimate, as it relies on specific autocorrelation estimates, which can be biased according to our simulations. Although negative genomic autocorrelation leads to reduced heritability compared to this commonly used definition, negative autocorrelation can also lead underestimation of genetic effects due to linkage masking. As these biases act in opposite directions it has been suggested that they might cancel each other out^14^, though the overall impact of genomic autocorrelation on these heritability estimation methods has not been explicitly analyzed.

## Discussion

We developed a new method, ACLR, to estimate genomic autocovariance and autocorrelation of causal effects on human complex traits as a function of base pair distance. We determined that ACLR is unbiased with well-calibrated standard errors in null simulations with zero autocorrelation; ACLR produced biased autocorrelation estimates in causal simulations with nonzero autocorrelation under some genetic architectures, but always detected the correct sign of the autocorrelation. Thus, any significantly nonzero autocorrelation reported by ACLR is a robust result, in the sense that the true autocorrelation must be nonzero with the reported sign. We applied ACLR to 31 diseases and complex traits from the UK Biobank^24^, detecting significantly negative autocorrelation at very short genomic distance ranges (up to 100bp) when meta-analyzing results across traits. The negative autocorrelation could be explained by opposite trait effects of linked SNPs cancelling each other out and escaping the action of negative selection, a phenomenon known as linkage masking^19^. The very short genomic distance range of the negative autocorrelation is unsurprising, as linkage masking effects at very short genomic distance range could be amplified by functional similarity at this range, e.g. due to compensatory mutations in binding sites or coding regions; SNP with opposite effects may be much less likely to arise at longer ranges. We further determined that the negative autocorrelation is primarily driven by pairs of SNPs in positive LD, consistent with the linkage masking hypothesis. We note that the negative autocorrelation has significant implications for association mapping, heritability estimation, and genetic risk prediction (see below).

Several previous studies have provided specific examples of opposite (and concordant) trait effects in humans and model organisms (although none of these studies estimated genome-wide autocorrelation of causal trait effects). In humans, Brown et al. ^19^ used a multi-SNP association approach to identify linked pairs of SNPs with opposite effects, on the other hand results by Zhou et al.^20^ based on deep learning-predicted genetic effects showed that mutational effects on gene expression often tend to be concordant within genes. In model organisms, Bernstein et al. ^23^ used high throughput phenotyping in *Caenorhabditis elegans* to identify several instances in which neighboring genetic variants have opposite effects, and She and Jarosz ^22^ estimated single nucleotide effects of several quantitative traits in a yeast cross experiment, finding an excess of nearby mutations that are functionally coupled, with examples of both aligned and opposite signed effects. Brown et al. ^19^ attributed their findings to linkage masking caused by negative selection; however, our results imply that linkage masking is far more pervasive than previously known. (We note that Brown et al.^19^ introduced the term linkage masking to emphasize linked genetic variants with opposite signed effects escaping detection in GWAS rather than escaping natural selection). Negative selection is known to broadly impact complex trait architectures, resulting in MAF-dependent^4;9;15;33^, LD-dependent^8^ and extremely polygenic^14^ architectures. Recently, Garcia and Lohmueller^21^ showed that negative autocorrelation of fitness effects of linked SNPs can arise under additive negative selection in evolutionary forward simulations, and detected signatures of this effect when comparing synonymous and non-synonymous SNP pairs in 1000 Genomes Project data^36^; however, Garcia and Lohmueller ^21^ did not analyze complex trait data. In addition, Zhou et al. ^37^ developed a method to estimate covariances between complex trait effects across fixed sets of genetic variants; however, unlike the method presented here, their method cannot be used to estimate autocovariance or autocorrelation.

Our results have several implications for future work. First, they indicate that linked SNPs with opposite effects are pervasive throughout the genome, likely leading to many disease risk variants that are undetected (or detected with underestimated effect) in GWAS. This motivates the use of improved association methods that allow for joint testing of multiple proximal variants ^19^. Second, our findings violate the assumption of uncorrelated SNP effects commonly used in fine mapping methods ^38^. Correcting this assumption might substantially improve the accuracy of these methods. Third, our results have important implications for defining and estimating SNP-heritability^2–6;8;10;12–15^. We show that negative causal minor allele effect autocorrelation between linked SNPs may cause true SNP-heritability to be substantially smaller than the heritability defined under these models, although further work is needed to conclusively determine the impact of negative autocorrelation on SNP-heritability estimates. Fourth, negative autocorrelation may contribute to trans-ethnic disparities in polygenic risk prediction^39–41^, as SNPs with opposite causal effects that are linked in the training population but not the target population are a source of heritability in the target population that cannot be predicted using the training population.

We note several limitations in our work. First, ACLR is unbiased and well-calibrated when the true autocorrelation is zero, but can produce biased autocorrelation estimates when the true autocorrelation is nonzero, though it detects the correct sign. Second, ACLR’s correction for MAF- and LD-dependent genetic architectures is essential. Our correction is based on a MAF- and LD-dependent genetic architecture model that is supported by empirical data^8;33;42;43^, but we cannot guarantee that this model is perfectly accurate. Third, we have not partitioned genomic autocorrelations by functional annotation or allele frequency (of one or both SNPs). As functional annotations and allele frequency ranges are differentially impacted by negative selection^8^, we hypothesize that genomic autocorrelations are likely to differ, making this an interesting direction for future analyses. Fourth, we recommend that ACLR should be applied using in-sample LD (analogous to recent studies in other settings^27–29^), which may be unavailable for some data sets. Fifth, our analysis focuses on minor allele effects but does not consider derived allele effects. While minor and derived alleles mostly match for SNPs with lower MAF, this is not the case for very common minor alleles. We note that our heritability shrinkage analysis is not affected by this definition, since switching the reference allele changes the sign of both LD and effect concordance. Sixth, our method does not model nonzero directional signed minor allele effects genome-wide, which might be expected for traits under directional selection. However, under our model, such effects would be expected to lead to positive long-range autocorrelation estimates, which we did not observe. Seventh, we did not include very rare SNPs (MAF < 0.1%) in our analysis. Shared tagging of unmodeled, very rare causal SNPs could produce spurious positive genomic autocorrelations, but would not be expected to lead to the negative genomic autocorrelations that we report. Finally, analogous to other studies that employ linear complex trait models, we have not investigated the potential impact of epistatic interactions on our estimates; however, the impact of epistatic interaction on these models is hypothesized to be small^44;45^. We note that epistatic interactions have also been suggested to alter selection effects on nearby genetic variants^23;46^ and might hence modulate the effect of linkage masking. Despite these limitations, our work provides an improved understanding of genomic autocorrelations of causal minor allele effect sizes.

## Acknowledgements

We are grateful to Evan Koch, Benjamin Neale, David Reich, Shamil Sunyaev, and Martin Zhang for helpful discussions. This research was conducted using the UK Biobank Resource under Application Number 16549 and was funded by NIH grants R01 MH101244, R37 MH107649 and U01 HG009088. Computational analyses were performed on the Orchestra High-Performance Compute Cluster at Harvard Medical School.

## Methods

### approach of the ACLR method

We assume a complex trait model, where genetic variants in the genome have a linear effect on the target trait and the trait is also affected by non-genetic environmental noise:

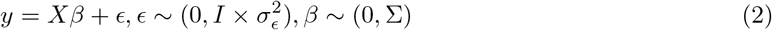

Here, *y* is the mean-centered, variance standardized trait vector, *X* is the column-mean-centered genotype matrix (minor allele counts with column means set to zero), each row representing one study individual and each column a genetic variant locus. ϵ is the environmental effect vector, which we assume to be uncorrelated between individuals. Furthermore, we define *β* as the vector of causal minor allele effects of genetic variants across the genome. In our model, these effects are random, drawn from an unspecified distribution with mean zero and covariance matrix Σ. Note that we do not constrain off-diagonal elements in Σ to be zero, which allows correlated minor allele effects. Since certain elements in *β* might be zero or negligibly small, this model does not necessarily assume that all variant loci in the genome have significant effects on the target trait.

We define summary statistics of variant *i* as 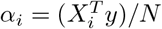 and in-sample LD matrix *R* = (*X^T^X*)/*N*, with *N* representing the number of individuals in the data set. From Equation 2 follows

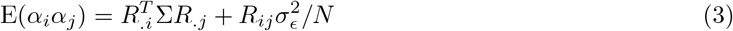

with *R_.i_* being the *i*^th^ column of *R*. Intuitively, this equation states that the correlation of summary statistics is a function of both the LD matrix as well as the correlation structure of the causal genetic effects, plus a term related to the environmental noise.

A critical assumption in our model is that the covariance between a pair of genetic effects is a function of the base pair distance between them, i.e. Σ_*ij*_ is a function of the distance between locus *i* and locus *j*. Specifically, we define *n* distance ranges with constant covariances, with the covariance between effects of variants *i* and *j* being *σ_k_* if the distance between the two loci falls into the *k^th^* distance range. In this case we can decompose Σ into a sum of distance dependent components:

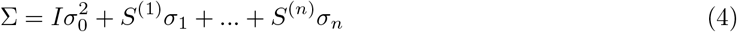

Here, *S*^(*k*)^ the design matrix of distance range *k*, where 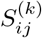 is *i* if the distance between variant *i* and *j* falls into the *k*^th^ distance range, and 0 otherwise. Combining Equations 3 and 4, we get

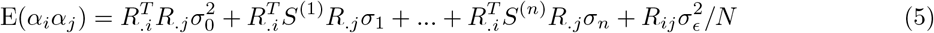

Distance dependent genetic effect covariances *σ_k_* can then be estimated by regressing the product of summary statistics *α_i_,α_j_* jointly on distance dependent LD scores 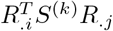. If model assumptions hold, this method will produce unbiased estimates of *σ_k_*.

### Details of ACLR and implementation

We developed ACLR, a software package implementing the above inference procedure. UK Biobank genotype data is read in from the compressed binary file format BGEN (see URLs) using a fast costumery C++ subroutine. For computational feasibility, in-sample LD matrices are calculated with a banded matrix approach, specifically we assume that LD between variant sites more than 1Mb apart are zero. We developed a Python implementation based on sparse matrix methods to efficiently calculate distance-dependent LD scores as defined above. Summary statistics were either calculated following the above definition of α¿ or using the corresponding mixed model association summary statistic calculated using BOLT-LMM^7^ to increase statistical power.

We used a weighted regression when estimating effect covariances to further improve power. Following the derivation of Bulik-Sullivan et al. ^26^, regression weights were based on the variance of the regressand under a normal model, specifically 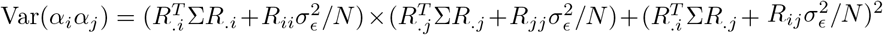. Since Σ and 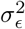 are unknown prior to inference, we assumed a diagonal covariance matrix with diagonal entries and 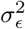 corresponding to a 50% heritable trait. Note that approximations of the variance of the regressand used to calculate regression weights cannot bias estimates but might only lead to suboptimal statistical power. Correlation results reported in the analysis are defined as 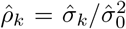 (see Equation 5). Uncertainty in 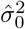 and 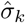 was inferred using a block jackknife approach, dividing the genome into 200 equally sized blocks of variants (10 jackknife blocks were used in simulations based on chromosome 22 only). The reported uncertainty in 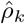 is based on the assumption of 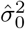 and 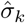 being independent and normally distributed. If model assumptions hold, 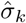 are unbiased estimates of *σ_k_* independent of the set of SNP pair summary statistics used, however they do affect statistical power. Maximum base pair distances between summary statistics pairs were considered in simulations, showing comparable power when only using *α_i_α_j_* terms with *i* = *j*. We therefore only used these terms in the analyses shown in this work after consideration these simulation results and computational constraints, even though ACLR allows using *i* ≠ *j* cross terms too.

Equation 4 implicitly assumes that all SNPs have the same effect size variance 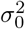. However, if SNPs with certain MAF and LD patterns have consistently larger or smaller effects, this assumption is broken and our method can yield biased results, consistent with previous work from related methods^8^. We devised a MAF- and LD-corrected version of ACLR that allows for effect size variance differences depending on the MAF and LD of SNPs and can remove this bias. Allowing for these differences assures that the model stays well specified in the presence of MAF- and LD-dependent architectures and hence mitigates possible biases in SNP effect autocovariance estimates. Following a model by Gazal et al.^33^, we divided all SNPs into 10 MAF bins for common SNPs (MAF ≥ 5%) and 10 MAF bins for low-frequency SNPs (MAF < 5%), for each choosing the MAF thresholds such that an equal number SNPs falls in each one of them. We then allow SNPs in each bin to have a distinct SNP effect variance. Furthermore, we used 6 LD-related functional annotations from Gazal et al.^33^, specifically predicted allelic age, recombination rate, level of LD in African populations, nucleotide diversity, background selection statistic, and CpG content. For all these LD-related annotations, apart from allelic age, we allow for different effect coefficients for low-frequency and common SNPs. These annotations linearly affected the SNP effect variances, following previous models^5;8^. Specifically, Equation 4 is replaced by

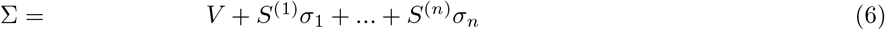

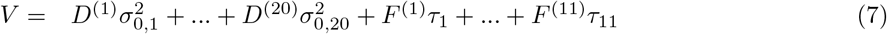

where *V* is the diagonal matrix of SNP effect variances, 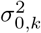 is the effect variance of the *k*^th^ MAF bin and *D*^(*k*)^ is a diagonal matrix with *D*^(*k*)^ being 1 if the *i*^th^ SNP falls into the *k*^th^ MAF bin and 0 otherwise. *F*^(*k*)^ is a diagonal matrix with 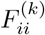 being the value of the *k*^th^ LD annotation of SNP *i* and *τ_k_* the linear effect of the annotation on the SNP effect variance. Equation 5 is then replaced by

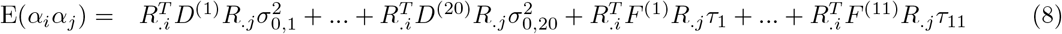

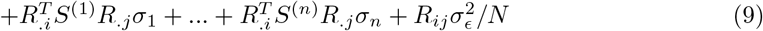

Covariances *σ_k_* are calculated by jointly solving this linear regression model for all variance and covariance parameters 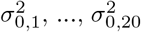, *τ*_1_,…, *τ*_11_, *σ*_1_,…, *σ_n_* and 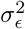. Autocorrelation estimates 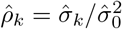 were calculated using the covariance estimates 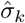 from the MAF-LD corrected method, but 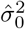 from the uncorrected method, since the former does not directly yield an estimate of the overall SNP effect variance and using an appropriate combination 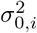 and *τ_i_* did not improve the accuracy of the autocorrelation estimate.

### UK Biobank genotype data

In our analysis we used genotype data from UK Biobank data set (see URLs). To avoid problems with population stratification, we only used data from 337, 502 individuals with self-reported and confirmed British ancestry that are unrelated (pairwise genetic relatedness < 5% after LD-pruning). We used genotype information of autosomal variant loci that were imputed using sequencing data from the UK10K project^47^ and 1000 Genomes Phase 3 (ref.^36^). These variant sites include both SNPs and small insertions and deletions, but for simplicity we are referring to all of them as SNPs. We only used SNPs with MAF > 0.1% to avoid using poorly imputed rare variants, resulting in 16, 525, 661 variant loci genome wide. Genotype data was read in from BGEN format files (see URLs), which contains the posterior probabilities after imputation of a given locus being homozygous reference, homozygous alternative and heterozygous, for each individual in the data set. In the analysis performed here, we used minor allele dosage genotypes (i.e. the expected number of minor alleles given these probabilities) as genotype values. Summary statistics were only calculated for a subset of SNPs, specifically SNPs from the HapMap Project Phase 3 (ref. ^48^). However, we did not use summary statistics from SNPs close to the human leukocyte antigen genes, specifically SNPs between 25.5 and 33.5Mb into chromosome 6.

### Simulations

All simulations used genotypes from 337,502 unrelated UK Biobank individuals with self-reported and confirmed British ancestry (see above section). Due to the large number of simulations, we restricted most simulations to chromosome 22 (232, 046 UK Biobank SNPs with MAF ≥ 0.1%); however, we also performed a selected set of simulations using genome-wide data (16, 525, 661 UK Biobank SNPs with MAF ≥ 0.1%; see above section). For null simulations, i.e. simulations with uncorrelated SNP effects, SNP effects *β* were drawn from independent mean-zero normal distributions. Depending on the specific simulation analysis, the effect variances were either identical for all SNPs, or varied according the inferred variances by Gazal et al.^33^, using their MAF and LD related annotations mentioned above. Since the model by Gazal et al. ^33^ infers effect variances for variance-standardized genotypes, in contrast to the non-standardized genotypes in this analysis, we rescaled MAF bin variances and LD-related variance effects by 2*p*(1 – *p*), where *p* is the median MAF of the respective MAF range. SNPs that had negative effect variances under this model, were assigned zero trait effects. Although we did not explicitly fix trait polygenicity in simulations, polygenicity is implicit in simulations that use MAF and LD related annotations, as they result in 27% of SNPs having negative effect variances and hence zero trait effects.

For causal simulations, correlated SNP effects were created by dividing all SNPs into blocks of *n* = 3 adjacent SNPs. In each block, SNP effects were drawn from a multivariate normal distribution with pairwise effect correlation being set to a fixed correlation parameter *ρ*_block_ = ±0.4. Effects of SNPs from different blocks were independent. Note that by construction, *ρ*_block_ does not correspond to the true correlation for any base-pair distance range used in the inference method. Instead, we used the sample correlation of all SNP pairs in a given distance bin as the ground truth to compare our inference results to. When using a MAF and LD-dependent architecture, effect variances were set as described for the null simulations above.

By construction, the above described simulations do not have constant effect covariances per distance range. We also performed causal simulations where we kept the effect covariance constant within each distance range. For simulations without MAF- and LD-dependent (and hence constant) effect variances, SNP base pair positions where altered in simulations such that blocks of *n* = 3 SNPs were within 100bp of each other, while other SNPs were further away, guaranteeing a fixed effect covariance within these blocks and zero covariance between blocks. In simulations with MAF- and LD-dependent variances and constant covariances per distance range, we chose a simpler architecture model than the one by Gazal et al. ^33^, which makes it possible to have constant covariances per distance range, without effect covariance matrices being not positive semi-definite. Specifically, effect variances were set to 1.0 for low-frequency SNPs (< 5% MAF) and 0.6 for common SNPs (> 5% MAF). Also, we chose the effect of the LLD African functional annotation^33^ to be τ = —0.1 and zero for all other annotations.

Given a set of genome-wide SNP effects that were simulated, trait values were then simulated using Equation 2. 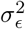 was chosen such that for a given heritability value, 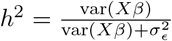.

### UK Biobank phenotype data

We analyzed 31 heritable traits and diseases from the UK Biobank data set (see URLs). Specifically, for each trait included in the final analysis we required heritability estimates from our method to be at least 6 standard errors away from zero. Furthermore, we removed all traits with missing trait values for more than 10% of the 337, 502 individuals used in this study for the following reason: the distance-dependent LD scores were only calculated once for all traits due to the large computational cost of this step; since our method assumes that summary statistics and LD scores are computed on the same set of individuals, we only included traits where this is approximately correct. The resulting set of 31 traits and diseases includes 9 quantitative traits, 13 case-control traits, as well as 9 quantitative blood cell traits (see Supplementary Table 2). When calculating summary statistics using the BOLT-LMM software^7^, we included sex, age, age squared, the assessment center, the genotyping platform, and 20 top principal components of the genotype matrix as fixed effects to mitigate possible confounding.

### Stratifying pairs of SNPs by their pairwise LD

A basic assumption of our method is that the SNP effect covariance only depends on the base pair distance between the respective SNP pair. However, we also performed an analysis that additionally stratified SNP pairs by pairwise LD between them. Specifically, instead of Equation 4, we now assume

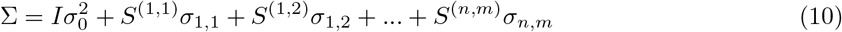

where *σ_i,j_* is the covariance of effects between SNP pairs that fall into the *i*^th^ distance range and the *j*^th^ pairwise LD bin, with *S*^(*i,j*)^ being defined accordingly. Here, pairwise LD is defined as the Pearson correlation coefficient *r*_LD_ between the minor allele counts of two SNPs in the study population. We define four pairwise LD bins: negative LD (−1 ≤ *r*_LD_ < −0.01), no LD (−0.01 ≤ *r*_LD_ < 0.01), moderate positive LD (0.01 ≤ *r*_LD_ < 0.3), strong positive LD (0.3 ≤ *r*_LD_ ≤ 1). For computational feasibility we reduced the distance bin number to three in these analyses: 1-100bp, 100-1000bp, >1000bp. Hence, *σ*_2,3_ here defines the effect covariance of SNP pairs that are 100-1000bp apart and the correlation between their respective minor alleles is between 0.01 and 0.3.

Estimation of *σ_ij_* parameters was performed identically to purely distance dependent covariance parameters and included MAF and LD bias correction, i.e. allowing for MAF and LD dependent SNP effect variances. For computational feasibility, LD between SNP pairs defining *S*^(*i,j*)^ were calculated only using a random subset of 1000 individuals.

### Heritability shrinkage estimation

In the special case of all SNP effects being uncorrelated and the trait variance standardized to 1, the linear complex trait model described in Equation 2 implies that the trait heritability is equal to the genome-wide sum of variances of standardized genotype effects, a fact that has been used in several previous heritability estimation methods^2–6;8;10;12–15^. We call this quantity 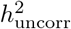, since it is only equal to the heritability if SNP effects are uncorrelated, but this is not true in the case of significant effect autocorrelation. If linked SNPs have systematic negatively correlated allele effects, these effects will cancel each other out and the heritability will be smaller than the sum of squared standardized SNP effects. Given a general linear complex trait model that allows correlated allele effects, we derive the following extended definition of heritability:

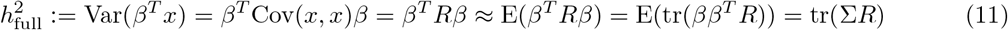

Similar to Equation 2, *R* is the genome-wide population LD matrix which we define as the minor allele count covariance matrix, *x* is a random genotype (i.e. minor allele count; not variance standardized but mean centered) vector, *β* is the causal minor allele effect vector, and Σ is the covariance matrix of *β*. The approximation is accurate as long as the trait is sufficiently polygenic. The equality tr(Σ*R*) = ∑_*i,j*_ (Σ ∘ *R*)_*i,j*_, the grand sum of the element-wise product, directly shows how positive LD anticorrelated effect SNPs as well as negative LD correlated effect SNPs reduce heritability. This is in contrast to the aforementioned definition used in previous analyses which assume uncorrelated effects, in which case Σ is a diagonal matrix and hence 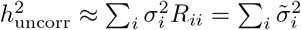, where 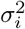 is the variance of the *i^th^* per-allele effect and 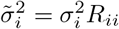 the variance of the *i^th^* allele effect after standardizing the genotype variance.

In order to assess the genome-wide magnitude of the cancelling of linked anticorrelated effects, we defined a new measure called ”heritability shrinkage” equal to 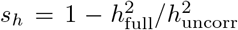, the fraction the heritability is reduced due to correlated SNP effects. We use our meta-analyzed UK Biobank trait autocorrelation estimates to calculate a rough estimate of *s_h_*, to get an approximate quantification of the impact of effect autocorrelation genome-wide. To do that, we ignore the MAF- and LD-dependent genetic architecture and assume 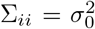 for all SNPs in the genome. Following our pairwise LD and distance stratification model of Σ in Equation 10, it follows that

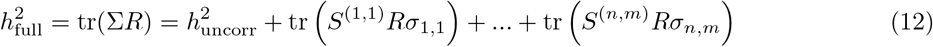

and hence,

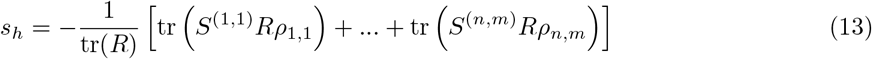

In practice, we calculated *s_h_* following the above equation using our pairwise LD and distance stratified autocorrelation estimates in Supplementary Table 9 and a banded LD matrix estimated from chromosome 10. Standard errors are directly based on the uncertainties in 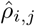 in Supplementary Table 9.

## URLs

Open-source software package implementing ACLR, https://github.com/arminschoech/ACLR; UK Biobank website, http://www.ukbiobank.ac.uk/; BGEN file format, http://www.well.ox.ac.uk/~gav/bgen_format/; UK Biobank genotype imputation manual, http://www.ukbiobank.ac.uk/wp-content/uploads/2014/04/imputation_documentation_May2015.pdf;

## Notes

### Competing Interest Statement

The authors have declared no competing interest.

