## Supplementary Information for "Negative short-range genomic autocorrelation of causal effects on human complex traits"

Schoech et al.

**Supplementary Table 1:** Whole-genome null simulations

| MAF-LD architect | MAF-LD corrected | $\hat{\rho}_{1-100bp}$ | $\hat{\rho}_{100-1000bp}$ | $\hat{\rho}_{>1000bp}$ |
|---|---|---|---|---|
| yes | no | $0.16 \pm 0.01$ | $-0.18 \pm 0.00$ | $0.01 \pm 0.00$ |
| yes | yes | $-0.00 \pm 0.01$ | $-0.01 \pm 0.00$ | $0.00 \pm 0.00$ |

Results from applying our method to 100 simulated traits based on genetic effects from all 22 autosomal chromosomes. Mean estimates and standard errors in the mean are shown. True correlation is zero in all simulations and distance ranges. Traits are simulated using MAF- and LD-dependent effect variances, using our method with and without MAF- and LD-correction.

**Supplementary Table 2:** Sample sizes and heritability estimates for 31 UK Biobank traits

| trait | sample size | $\hat{h}^2$ |
|---:|:---:|:---:|
| blood pressure (diastolic) | 310,831 | 0.256±0.013 |
| blood pressure (systolic) | 310,831 | 0.262±0.013 |
| body mass index | 336,394 | 0.308±0.014 |
| bone mineral density | 327,738 | 0.390±0.039 |
| cholesterol | 321,595 | 0.167±0.024 |
| height | 336,759 | 0.556±0.037 |
| low-density lipoprotein | 321,002 | 0.089±0.014 |
| smoking status | 336,312 | 0.057±0.003 |
| waist-hip ratio | 336,847 | 0.193±0.013 |
| AID disease | 337,488 | 0.005±0.001 |
| asthma | 337,071 | 0.008±0.001 |
| cardiovascular disease | 337,488 | 0.028±0.001 |
| college education | 334,353 | 0.040±0.002 |
| diabetes (any) | 336,759 | 0.002±0.000 |
| diabetes (type 2) | 337,488 | 0.002±0.000 |
| diabetes (endocrine) | 337,488 | 0.002±0.000 |
| eczema | 337,071 | 0.016±0.002 |
| high cholesterol | 337,488 | 0.005±0.001 |
| hypertension | 336,972 | 0.032±0.002 |
| hypothyroidism | 337,488 | 0.003±0.000 |
| respiratory disease | 337,488 | 0.009±0.001 |
| thyroid disease | 337,488 | 0.003±0.000 |
| eosinophil count | 323,358 | 0.280±0.032 |
| high light scatter reticulocyte count | 321,608 | 0.262±0.032 |
| lymphocyte count | 326,304 | 0.244±0.016 |
| mean corpuscular hemoglobin | 326,188 | 0.358±0.047 |
| monocyte count | 325,319 | 0.275±0.035 |
| platelet count | 326,621 | 0.413±0.040 |
| red blood cell count | 327,209 | 0.272±0.031 |
| red blood cell distribution width | 325,395 | 0.303±0.031 |
| white blood cell count | 326,723 | 0.247±0.020 |

This table shows the number of phenotype values used as well as heritability estimates corresponding to SNP effect variance estimates $\hat{\sigma}_0^2$ as estimated by ACLR (see Methods) for 9 quantitative traits, 13 case-control traits, and 9 quantitative blood cell traits. Specifically, we used the definition of heritability $\hat{h}^2 = \sum_i R_{ii}\hat{\sigma}_0^2$ over all SNPs genome-wide, where $R$ is the LD matrix, specifically the minor allele count in-sample covariance matrix. Note that we did not account for heritability shrinkage in this analysis (see last paragraph of Results). Also note that the table shows observed-scale heritability values for case-control traits (see Lee et al. 2011 AJHG).

**Supplementary Table 3:** Autocorrelation estimates for 31 UK Biobank traits

| trait | $\hat{\rho}_{1-50bp}$ | $\hat{\rho}_{50-100bp}$ | $\hat{\rho}_{100-200bp}$ | $\hat{\rho}_{200-500bp}$ | $\hat{\rho}_{500-1000bp}$ |
|---|---|---|---|---|---|
| blood pressure (diastolic) | -0.52 $\pm$ 0.19 | -0.31 $\pm$ 0.32 | -0.12 $\pm$ 0.19 | -0.02 $\pm$ 0.10 | 0.12 $\pm$ 0.06 |
| blood pressure (systolic) | -0.30 $\pm$ 0.18 | -0.27 $\pm$ 0.32 | -0.12 $\pm$ 0.21 | -0.03 $\pm$ 0.10 | 0.11 $\pm$ 0.07 |
| body mass index | -0.23 $\pm$ 0.22 | -0.53 $\pm$ 0.24 | 0.01 $\pm$ 0.20 | -0.09 $\pm$ 0.11 | 0.18 $\pm$ 0.09 |
| bone mineral density | -0.18 $\pm$ 0.53 | -0.09 $\pm$ 0.54 | 0.49 $\pm$ 0.55 | -0.18 $\pm$ 0.38 | 0.06 $\pm$ 0.26 |
| cholesterol | -0.86 $\pm$ 0.99 | -1.39 $\pm$ 0.95 | -1.44 $\pm$ 1.04 | 0.35 $\pm$ 0.45 | 0.30 $\pm$ 0.29 |
| height | -0.40 $\pm$ 0.21 | -0.45 $\pm$ 0.26 | 0.09 $\pm$ 0.18 | 0.03 $\pm$ 0.09 | 0.05 $\pm$ 0.06 |
| low-density lipoprotein | -2.01 $\pm$ 1.42 | -1.67 $\pm$ 1.27 | -0.51 $\pm$ 0.81 | 0.38 $\pm$ 0.61 | 0.26 $\pm$ 0.35 |
| smoking status | -0.44 $\pm$ 0.23 | -0.30 $\pm$ 0.31 | -0.25 $\pm$ 0.22 | 0.17 $\pm$ 0.11 | 0.07 $\pm$ 0.06 |
| waist-hip ratio | -0.51 $\pm$ 0.29 | -0.27 $\pm$ 0.38 | -0.42 $\pm$ 0.25 | -0.11 $\pm$ 0.14 | 0.25 $\pm$ 0.07 |
| AID disease | 0.20 $\pm$ 0.75 | -0.94 $\pm$ 0.96 | 1.10 $\pm$ 0.58 | -0.18 $\pm$ 0.31 | -0.07 $\pm$ 0.21 |
| asthma | -0.49 $\pm$ 0.45 | -0.14 $\pm$ 0.59 | 0.25 $\pm$ 0.37 | 0.06 $\pm$ 0.17 | -0.05 $\pm$ 0.10 |
| cardiovascular disease | -0.29 $\pm$ 0.21 | 0.15 $\pm$ 0.35 | -0.07 $\pm$ 0.24 | -0.06 $\pm$ 0.12 | 0.07 $\pm$ 0.08 |
| college education | -0.40 $\pm$ 0.17 | -0.18 $\pm$ 0.23 | -0.36 $\pm$ 0.14 | 0.12 $\pm$ 0.09 | 0.06 $\pm$ 0.06 |
| diabetes (any) | 0.18 $\pm$ 0.59 | -0.02 $\pm$ 0.93 | -0.59 $\pm$ 0.54 | -0.08 $\pm$ 0.27 | 0.32 $\pm$ 0.26 |
| diabetes (type 2) | 0.14 $\pm$ 0.60 | -0.38 $\pm$ 0.94 | -0.66 $\pm$ 0.56 | -0.13 $\pm$ 0.27 | 0.43 $\pm$ 0.27 |
| diabetes (endocrine) | 0.32 $\pm$ 0.67 | -0.00 $\pm$ 0.98 | -0.35 $\pm$ 0.64 | -0.30 $\pm$ 0.32 | 0.40 $\pm$ 0.27 |
| eczema | 0.11 $\pm$ 0.56 | -0.79 $\pm$ 0.47 | -0.15 $\pm$ 0.30 | -0.04 $\pm$ 0.16 | 0.20 $\pm$ 0.11 |
| high cholesterol | -0.18 $\pm$ 0.61 | 0.15 $\pm$ 0.87 | -0.74 $\pm$ 0.74 | -0.03 $\pm$ 0.46 | 0.25 $\pm$ 0.24 |
| hypertension | -0.32 $\pm$ 0.21 | 0.01 $\pm$ 0.32 | -0.13 $\pm$ 0.22 | 0.04 $\pm$ 0.11 | 0.02 $\pm$ 0.08 |
| hypothyroidism | 0.39 $\pm$ 0.85 | -0.49 $\pm$ 0.99 | 0.89 $\pm$ 0.58 | -0.26 $\pm$ 0.33 | -0.01 $\pm$ 0.25 |
| respiratory disease | -0.74 $\pm$ 0.44 | -0.07 $\pm$ 0.61 | 0.47 $\pm$ 0.40 | 0.01 $\pm$ 0.19 | -0.03 $\pm$ 0.10 |
| thyroid disease | 0.64 $\pm$ 0.93 | -1.00 $\pm$ 1.11 | 1.05 $\pm$ 0.59 | -0.22 $\pm$ 0.34 | -0.07 $\pm$ 0.25 |
| eosinophil count | -0.64 $\pm$ 0.32 | -0.60 $\pm$ 0.41 | 0.12 $\pm$ 0.31 | -0.05 $\pm$ 0.16 | 0.08 $\pm$ 0.11 |
| high light scatter reticulocyte count | 0.71 $\pm$ 0.82 | 0.85 $\pm$ 1.50 | -0.29 $\pm$ 0.61 | -0.32 $\pm$ 0.30 | 0.10 $\pm$ 0.10 |
| lymphocyte count | 0.28 $\pm$ 0.45 | -0.11 $\pm$ 0.46 | -0.60 $\pm$ 0.44 | -0.20 $\pm$ 0.19 | 0.25 $\pm$ 0.14 |
| mean corpuscular hemoglobin | -0.03 $\pm$ 0.53 | -1.56 $\pm$ 1.81 | -0.19 $\pm$ 0.87 | 0.83 $\pm$ 0.40 | -0.31 $\pm$ 0.25 |
| monocyte count | 0.20 $\pm$ 0.59 | -1.53 $\pm$ 0.99 | -0.30 $\pm$ 0.45 | -0.06 $\pm$ 0.31 | 0.19 $\pm$ 0.25 |
| platelet count | -0.54 $\pm$ 0.26 | -0.42 $\pm$ 0.46 | 0.35 $\pm$ 0.37 | -0.01 $\pm$ 0.21 | 0.07 $\pm$ 0.13 |
| red blood cell count | -0.35 $\pm$ 0.35 | -0.70 $\pm$ 0.52 | -0.53 $\pm$ 0.46 | 0.26 $\pm$ 0.22 | 0.06 $\pm$ 0.15 |
| red blood cell distribution width | -0.35 $\pm$ 0.48 | -0.84 $\pm$ 1.29 | 0.07 $\pm$ 0.64 | 0.12 $\pm$ 0.32 | 0.15 $\pm$ 0.19 |
| white blood cell count | 0.42 $\pm$ 0.59 | -0.96 $\pm$ 0.51 | -0.05 $\pm$ 0.24 | -0.21 $\pm$ 0.14 | 0.21 $\pm$ 0.08 |
| meta-analysis | $-0.349 \pm 0.078$ | $-0.329 \pm 0.106$ | $-0.044 \pm 0.070$ | $-0.001 \pm 0.038$ | $0.071 \pm 0.024$ |

Estimated distance-dependent autocorrelation results are reported for 31 complex traits in the UK Biobank data set. These values correspond to results displayed in Figure 1. Estimates are reported with block-jackknife standard errors.

**Supplementary Table 4:** Autocovariance estimates for 31 UK Biobank traits

a) Distance ranges < 200bp

| trait | $\hat{\sigma}_{1-50bp}$ | $\hat{\sigma}_{50-100bp}$ | $\hat{\sigma}_{100-200bp}$ |
|---:|---:|---:|---:|
| blood pressure (diastolic) | -5.42e-8±2.01e-8 | -3.28e-8±3.34e-8 | -1.28e-8±1.95e-8 |
| blood pressure (systolic) | -3.17e-8±1.97e-8 | -2.88e-8±3.44e-8 | -1.31e-8±2.21e-8 |
| body mass index | -2.93e-8±2.79e-8 | -6.7e-8±2.96e-8 | 1.03e-9±2.54e-8 |
| bone mineral density | -2.84e-8±8.51e-8 | -1.45e-8±8.54e-8 | 7.85e-8±8.72e-8 |
| cholesterol | -5.89e-8±6.73e-8 | -9.53e-8±6.35e-8 | -9.85e-8±6.99e-8 |
| height | -9.02e-8±4.77e-8 | -1.01e-7±5.93e-8 | 2.06e-8±4.14e-8 |
| low-density lipoprotein | -7.29e-8±5.03e-8 | -6.05e-8±4.52e-8 | -1.83e-8±2.94e-8 |
| smoking status | -1.04e-8±5.28e-9 | -7.05e-9±7.33e-9 | -5.85e-9±5.03e-9 |
| waist-hip ratio | -4.03e-8±2.24e-8 | -2.09e-8±3.01e-8 | -3.35e-8±1.98e-8 |
| AID disease | 3.8e-10±1.45e-9 | -1.8e-9±1.83e-9 | 2.11e-9±1.08e-9 |
| asthma | -1.63e-9±1.47e-9 | -4.77e-10±1.95e-9 | 8.32e-10±1.23e-9 |
| cardiovascular disease | -3.39e-9±2.45e-9 | 1.69e-9±4.07e-9 | -8.13e-10±2.79e-9 |
| college education | -6.41e-9±2.78e-9 | -2.85e-9±3.69e-9 | -5.87e-9±2.3e-9 |
| diabetes (any) | 1.71e-10±5.56e-10 | -1.48e-11±8.75e-10 | -5.57e-10±5.06e-10 |
| diabetes (type 2) | 1.14e-10±4.81e-10 | -3.04e-10±7.51e-10 | -5.27e-10±4.43e-10 |
| diabetes (endocrine) | 2.13e-10±4.4e-10 | -1.09e-13±6.43e-10 | -2.32e-10±4.22e-10 |
| eczema | 6.96e-10±3.64e-9 | -5.15e-9±3.04e-9 | -1.01e-9±1.93e-9 |
| high cholesterol | -3.97e-10±1.31e-9 | 3.19e-10±1.88e-9 | -1.6e-9±1.58e-9 |
| hypertension | -4.16e-9±2.74e-9 | 7.3e-11±4.2e-9 | -1.73e-9±2.86e-9 |
| hypothyroidism | 4.55e-10±9.99e-10 | -5.82e-10±1.17e-9 | 1.05e-9±6.68e-10 |
| respiratory disease | -2.62e-9±1.51e-9 | -2.61e-10±2.16e-9 | 1.64e-9±1.39e-9 |
| thyroid disease | 8.92e-10±1.28e-9 | -1.39e-9±1.53e-9 | 1.46e-9±7.93e-10 |
| eosinophil count | -7.35e-8±3.52e-8 | -6.93e-8±4.66e-8 | 1.34e-8±3.52e-8 |
| high light scatter reticulocyte count | 7.59e-8±8.7e-8 | 9.11e-8±1.6e-7 | -3.07e-8±6.5e-8 |
| lymphocyte count | 2.79e-8±4.53e-8 | -1.1e-8±4.58e-8 | -6.01e-8±4.38e-8 |
| mean corpuscular hemoglobin | -4.62e-9±7.7e-8 | -2.29e-7±2.64e-7 | -2.78e-8±1.28e-7 |
| monocyte count | 2.2e-8±6.66e-8 | -1.73e-7±1.09e-7 | -3.4e-8±5.01e-8 |
| platelet count | -9.1e-8±4.35e-8 | -7.03e-8±7.77e-8 | 5.88e-8±6.17e-8 |
| red blood cell count | -3.88e-8±5.35e-8 | -9.4e-8±1.44e-7 | 7.9e-9±7.15e-8 |
| red blood cell distribution width | -4.38e-8±4.37e-8 | -8.63e-8±6.4e-8 | -6.52e-8±5.63e-8 |
| white blood cell count | 4.19e-8±5.94e-8 | -9.7e-8±5.1e-8 | -5.14e-9±2.4e-8 |

b) Distance ranges > 200bp

| trait | $\hat{\sigma}_{200-500bp}$ | $\hat{\sigma}_{500-1000bp}$ | $\hat{\sigma}_{>1000bp}$ |
|---|---|---|---|
| blood pressure (diastolic) | -1.99e-9±1.07e-8 | 1.28e-8±6.76e-9 | 2.95e-12±1.34e-11 |
| blood pressure (systolic) | -3.29e-9±1.09e-8 | 1.14e-8±7.14e-9 | 9.11e-12±1.3e-11 |
| body mass index | -1.09e-8±1.34e-8 | 2.28e-8±1.08e-8 | -9.6e-12±1.35e-11 |
| bone mineral density | -2.87e-8±6.01e-8 | 9.02e-9±4.22e-8 | -4.75e-12±2.51e-11 |
| cholesterol | 2.37e-8±3.07e-8 | 2.04e-8±1.94e-8 | 3.7e-11±4.29e-11 |
| height | 6.55e-9±2.14e-8 | 1.15e-8±1.39e-8 | 4.67e-11±4.1e-11 |
| low-density lipoprotein | 1.38e-8±2.19e-8 | 9.61e-9±1.27e-8 | 1.77e-11±2.72e-11 |
| smoking status | 3.87e-9±2.57e-9 | 1.6e-9±1.3e-9 | 2.34e-12±2.37e-12 |
| waist-hip ratio | -8.57e-9±1.14e-8 | 1.98e-8±5.66e-9 | -2.15e-12±9.19e-12 |
| AID disease | -3.47e-10±6.01e-10 | -1.27e-10±4.03e-10 | 1.84e-13±5.12e-13 |
| asthma | 1.93e-10±5.68e-10 | -1.56e-10±3.41e-10 | 2.61e-13±5.87e-13 |
| cardiovascular disease | -7.2e-10±1.35e-9 | 8.53e-10±9.54e-10 | 7.69e-13±1.37e-12 |
| college education | 1.98e-9±1.46e-9 | 1.05e-9±9.15e-10 | 2.37e-13±1.51e-12 |
| diabetes (any) | -8.02e-11±2.56e-10 | 3.05e-10±2.44e-10 | 2.08e-13±2.29e-13 |
| diabetes (type 2) | -1.06e-10±2.15e-10 | 3.42e-10±2.12e-10 | 9.13e-14±1.99e-13 |
| diabetes (endocrine) | -1.97e-10±2.11e-10 | 2.65e-10±1.73e-10 | 4.24e-14±1.85e-13 |
| eczema | -2.55e-10±1.06e-9 | 1.29e-9±6.78e-10 | -5.93e-13±1.16e-12 |
| high cholesterol | -7.51e-11±9.91e-10 | 5.35e-10±5.1e-10 | 7.62e-13±9.78e-13 |
| hypertension | 5.58e-10±1.47e-9 | 2.36e-10±1.03e-9 | 2.14e-13±1.36e-12 |
| hypothyroidism | -3.01e-10±3.81e-10 | -1.71e-11±2.9e-10 | 5.8e-14±3.25e-13 |
| respiratory disease | 4.03e-11±6.59e-10 | -1.21e-10±3.6e-10 | 3.29e-13±5.91e-13 |
| thyroid disease | -3.05e-10±4.7e-10 | -9.32e-11±3.51e-10 | -6.12e-14±3.85e-13 |
| eosinophil count | -6.18e-9±1.85e-8 | 8.73e-9±1.26e-8 | 5.02e-11±5.26e-11 |
| high light scatter reticulocyte count | -3.45e-8±3.22e-8 | 1.12e-8±1.03e-8 | 2.23e-11±3.04e-11 |
| lymphocyte count | -2.03e-8±1.91e-8 | 2.49e-8±1.4e-8 | 1.93e-11±2.52e-11 |
| mean corpuscular hemoglobin | 1.22e-7±5.64e-8 | -4.55e-8±3.63e-8 | -3.22e-11±2.27e-11 |
| monocyte count | -6.23e-9±3.49e-8 | 2.18e-8±2.78e-8 | -1.08e-11±1.41e-11 |
| platelet count | -2.0e-9±3.49e-8 | 1.13e-8±2.19e-8 | -3.67e-11±1.91e-11 |
| red blood cell count | 1.36e-8±3.58e-8 | 1.67e-8±2.1e-8 | -3.78e-11±1.98e-11 |
| red blood cell distribution width | 3.28e-8±2.69e-8 | 7.85e-9±1.83e-8 | 1.69e-12±1.65e-11 |
| white blood cell count | -2.16e-8±1.41e-8 | 2.12e-8±8.22e-9 | -4.51e-12±1.76e-11 |

Autocovariance estimates for 31 UK Biobank traits are shown with jackknife standard errors. These estimates correspond to the autocorrelation estimates in Figure 1, which are based on the autocovariance estimates shown here.

**Supplementary Table 5:** Testing for autocorrelation heterogeneity between traits

| | 1-50bp | 50-100bp | 100-200bp | 200-500bp | 500-1000bp | > 1000bp |
|---|---|---|---|---|---|---|
| P-value | 0.10 | 0.09 | 0.8 | 0.08 | 0.06 | 0.024 |

For each distance range P-values for heterogeneity were calculated based on the 31 autocorrelation estimates and standard errors in Figure 1. No distance range shows significant evidence for heterogeneity across traits, apart from the > 1000bp range which shows very small autocorrelation estimates (absolute value $< 10^{-3}$, see Supplementary Table 4) and is non-significant after correcting for six hypothesis tested. In each case we calculated the cross-trait best-fit autocorrelation as the inverse variance weighted mean estimate. Given this best-fit estimate and assuming that standard error estimates are sufficiently precise, we defined our null hypothesis as deviations from this best-fit estimate over the standard error estimate following a standard normal distribution. We then calculated P-values based on that assumption.

**Supplementary Table 6:** Autocorrelation estimates for the > 1000bp distance range

| trait | $\hat{\rho}_{>1000bp}$ |
|---|---|
| blood pressure (diastolic) | 2.81e-5±1.28e-4 |
| blood pressure (systolic) | 8.48e-5±1.21e-4 |
| body mass index | -7.61e-5±1.07e-4 |
| bone mineral density | -2.98e-5±1.57e-4 |
| cholesterol | 5.41e-4±6.31e-4 |
| height | 2.05e-4±1.81e-4 |
| low-density lipoprotein | 4.88e-4±7.54e-4 |
| smoking status | 1.00e-4±1.01e-4 |
| waist-hip ratio | -2.73e-5±1.17e-4 |
| AID disease | 9.6e-5±2.67e-4 |
| asthma | 7.89e-5±1.78e-4 |
| cardiovascular disease | 6.63e-5±1.18e-4 |
| college education | 1.46e-5±9.34e-5 |
| diabetes (any) | 2.19e-4±2.43e-4 |
| diabetes (type 2) | 1.14e-4±2.48e-4 |
| diabetes (endocrine) | 6.46e-5±2.82e-4 |
| eczema | -9.07e-5±1.78e-4 |
| high cholesterol | 3.54e-4±4.57e-4 |
| hypertension | 1.65e-5±1.05e-4 |
| hypothyroidism | 4.92e-5±2.76e-4 |
| respiratory disease | 9.35e-5±1.68e-4 |
| thyroid disease | -4.41e-5±2.78e-4 |
| eosinophil count | 4.37e-4±4.62e-4 |
| high light scatter reticulocyte count | 2.09e-4±2.85e-4 |
| lymphocyte count | 1.93e-4±2.53e-4 |
| mean corpuscular hemoglobin | -2.2e-4±1.58e-4 |
| monocyte count | -9.61e-5±1.26e-4 |
| platelet count | -2.17e-4±1.15e-4 |
| red blood cell count | -3.4e-4±1.82e-4 |
| red blood cell distribution width | 1.36e-5±1.33e-4 |
| white blood cell count | -4.46e-5±1.74e-4 |
| meta-analysis | 1.13e-05±3.82e-05 |

Autocorrelation estimates and standard errors for the > 1000bp distance range are shown for 31 UK Biobank traits like in the corresponding closer distance range results displayed in Figure 1. The meta-analysis was performed using a subset of 25 independent traits with genetic correlation less than 0.5. Following Equation 1, the results only represent SNP pairs that are connected to the summary statistic SNP by significant LD. Since our method assumes vanishing LD between SNPs more than 1Mb apart for computational reasons, this estimate only represents SNP pairs at most 2Mb apart, though is likely most representative of much shorter length scales, as LD is significantly reduced over the length scale of 1Mb.

**Supplementary Table 7:** Meta-analysis of autocorrelation estimates across 31 traits using different approaches

| MAF-LD corrected | $r^2_{\max}$ | $\hat{\rho}_{1-50bp}$ | $\hat{\rho}_{50-100bp}$ | $\hat{\rho}_{100-200bp}$ | $\hat{\rho}_{200-500bp}$ | $\hat{\rho}_{500-1000bp}$ |
|---|---|---|---|---|---|---|
| yes | 0.5 | -0.35 $\pm$ 0.08 | -0.33 $\pm$ 0.11 | -0.04 $\pm$ 0.07 | -0.00 $\pm$ 0.04 | 0.07 $\pm$ 0.02 |
| yes | 0.1 | -0.39 $\pm$ 0.10 | -0.32 $\pm$ 0.13 | -0.13 $\pm$ 0.09 | 0.04 $\pm$ 0.05 | 0.07 $\pm$ 0.03 |
| no | 0.5 | 0.15 $\pm$ 0.08 | -0.36 $\pm$ 0.11 | -0.19 $\pm$ 0.07 | -0.17 $\pm$ 0.04 | -0.08 $\pm$ 0.03 |
| no | 0.1 | 0.06 $\pm$ 0.10 | -0.36 $\pm$ 0.13 | -0.28 $\pm$ 0.09 | -0.12 $\pm$ 0.05 | -0.07 $\pm$ 0.03 |

Cross-trait meta-analyzed autocorrelation estimates using different variants of our method are shown. We include results for each of the five distance ranges with and without MAF- and LD-correction as well as at two different genetic correlation threshold ($r^2_{\max}$) to avoid genetically correlated traits biasing the results (see Methods). Out of the 31 UK Biobank traits analyzed, an $r^2_{\max}$ of 0.5 excludes 6, while an $r^2_{\max}$ of 0.1 excludes 18 traits due to high genetic correlation with another, more heritable trait.

**Supplementary Table 8:** Pairwise LD-stratified autocovariance inference in simulations

| $\sigma_{\text{1-100bp}}$ | $\hat{\sigma}_{\text{1-100bp,lowLD}}$ | $\hat{\sigma}_{\text{1-100bp,highLD}}$ | $\sigma_{\text{100-1000bp}}$ | $\hat{\sigma}_{\text{100-1000bp,lowLD}}$ | $\hat{\sigma}_{\text{100-1000bp,highLD}}$ |
|---|---|---|---|---|---|
| 0.0 | 0.021 $\pm$ 0.028 | 0.008 $\pm$ 0.021 | 0.0 | 0.004 $\pm$ 0.006 | -0.006 $\pm$ 0.004 |
| 0.2 | 0.204 $\pm$ 0.022 | 0.196 $\pm$ 0.016 | 0.0 | -0.006 $\pm$ 0.005 | -0.002 $\pm$ 0.003 |
| -0.2 | -0.166 $\pm$ 0.014 | -0.211 $\pm$ 0.010 | 0.0 | -0.001 $\pm$ 0.003 | -0.001 $\pm$ 0.002 |

Effect covariances used to simulate traits for each distance range together with mean estimates and standard errors in the mean are shown. Traits are simulated with MAF- and LD-dependent architectures and we used MAF- and LD-corrected ACLR. True effect covariances are constant within each of the two distance ranges, i.e. the same for all SNP pairs in a distance range independent of the LD between the pair of SNPs. We used the pairwise LD stratified extension of ACLR to test for differences between SNP pairs with different LD values. Low LD corresponds to SNP pairs that have no significant or negative LD (minor allele correlation $< 0.01$), while high LD indicates significant positive LD (minor allele correlation $\geq 0.01$). Results are based on 1000 simulations each, using SNPs from chromosome 22 only (MAF $\geq 0.1\%$)

**Supplementary Table 9:** Cross-trait meta-analysis of pairwise LD-stratified autocorrelations in UK Biobank

| distance bin | $r_{\text{LD}} < -0.01$ | $-0.01 \leq r_{\text{LD}} < 0.01$ | $0.01 \leq r_{\text{LD}} < 0.3$ | $0.3 < r_{\text{LD}}$ |
|---|---|---|---|---|
| 1-100bp | 0.256$\pm$0.087 | -1.652$\pm$2.038 | -0.279$\pm$0.389 | -0.180$\pm$0.035 |
| 100-1000bp | 0.038$\pm$0.012 | 0.588$\pm$0.321 | -0.227$\pm$0.053 | 0.003$\pm$0.006 |
| >1000bp | 0.000$\pm$0.000 | 0.000$\pm$0.000 | 0.000$\pm$0.000 | 0.000$\pm$0.000 |

Cross-trait autocorrelation estimates and standard errors when stratifying SNP pairs by pairwise LD additionally to base pair distance are displayed (see Methods). $r_{\text{LD}}$ corresponds to the LD value (Pearson correlation coefficient between minor allele counts) between SNP pairs. Cross-trait meta-analysis was performed using a subset of 25 independent UK Biobank with pairwise genetic correlation $< 0.5$, following the main analysis in Figure 1. Consistent with the linkage masking hypothesis, short-range (1-100bp) autocorrelations are significantly negative for SNPs in positive LD, but significantly positive for SNPs in negative LD. 100-1000bp autocorrelations exhibited a partially consistent pattern, despite the absence of a significant negative autocorrelation across all SNP pairs (see Figure 1). We note that positively correlated trait effects of alleles in negative LD could be explained by linkage masking, however positive autocorrelation of minor allele effects could also arise in conserved regions independent of local LD, as deleterious alleles will more frequently be minor.
